# nf-LO: A scalable, containerised workflow for genome-to-genome lift over

**DOI:** 10.1101/2021.05.25.445595

**Authors:** Andrea Talenti, James Prendergast

## Abstract

The increasing availability of new genome assemblies often comes with an impaired amount of associated genomic annotations, limiting the range of studies that can be performed. A common workaround is to lift over annotations from better annotated genomes. However, generating the files required to perform a liftover is computationally and labour intensive and only a limited number are currently publicly available.

Here we present nf-LO (nextflow-LiftOver), a containerised and scalable Nextflow pipeline that enables liftovers within and between any species for which assemblies are available. nf-LO will consequently facilitates data interpretation across a broad range of genomic studies.

## Main body

The advent of third generation sequencing and ultra-fast assemblers (Joseph et al. 2018; Ruan and Li 2020) allows for the generation of high quality *de novo* assemblies in a fraction of the previous time. As a result increasingly large numbers of new genomes for several species are being generated (Zoonomia consortium 2020).

Despite this increased availability, novel assemblies most often lack the extensive annotation data required to perform downstream analyses. Not only simple annotations such as gene models, but also supplementary resources for researcher to understand the biological significance of their studies. Unfortunately, such resources are generally only available for a small number of model organisms (OMIA; Amberger et al. 2015; Carithers and Moore 2015; Hu et al. 2019).

A solution to the problem is to liftover positions and annotation (i.e. cross-mapping of the loci) to the new genome from well-annotated assemblies, using tools such as LiftOver (Navarro Gonzalez et al. 2021) and NCBI Remap (Luu et al. 2020). However, the alignment files required to perform these analyses are only currently publicly available for a small number of pairs of genomes. For all other pairs of genomes researchers have to generate their own liftover files. Only a few algorithms address the problem in an easy to implement and distributable way, e.g. flo for same species liftovers (Pracana et al. 2017) and LiftOff for ultra-fast liftovers (Shumate and Salzberg 2020)). In this study we present nf-LO, a scalable workflow to generate liftover files for any pair of genomes based on the UCSC liftover pipeline. Nf-LO can directly pull genomes from public repositories, supports parallelised alignment using a range of alignment tools and can be finely tuned to achieve the desired sensitivity, speed of process and repeatability of analyses.

nf-LO is a workflow to facilitate the generation of genome alignment chain files compatible with the LiftOver utility. It is written in Nextflow, a domain specific language v2 (DSL) and workflow manager, that allows easy implementation, redistribution and scalability of complex workflows across every Unix-based operating system; ranging from a desktop machine to cloud computing and HPC clusters. The dependencies are shipped alongside the workflow as docker containers or as an anaconda environment, facilitating the diffusion and adoption of the workflow across different systems.

The software accepts any two input genomes in fasta format, or alternatively can download a resource by providing a web address, an iGenome identifier or an NCBI GenBank or RefSeq accession. The workflow is shown in Figure 1, and in brief consists of three core steps, and one optional one: 1) chunking the two genomes, 2) pairwise alignment of the blocks, 3) generating the chain-net file that can be used to perform the liftover and, if a bed/gff/gtf/vcf/bam/maf file is provided, 4) performing the liftover from source to target. The chunking approach dramatically reduces the runtime of the analysis by parallelizing the alignments.

**Figure 1.**
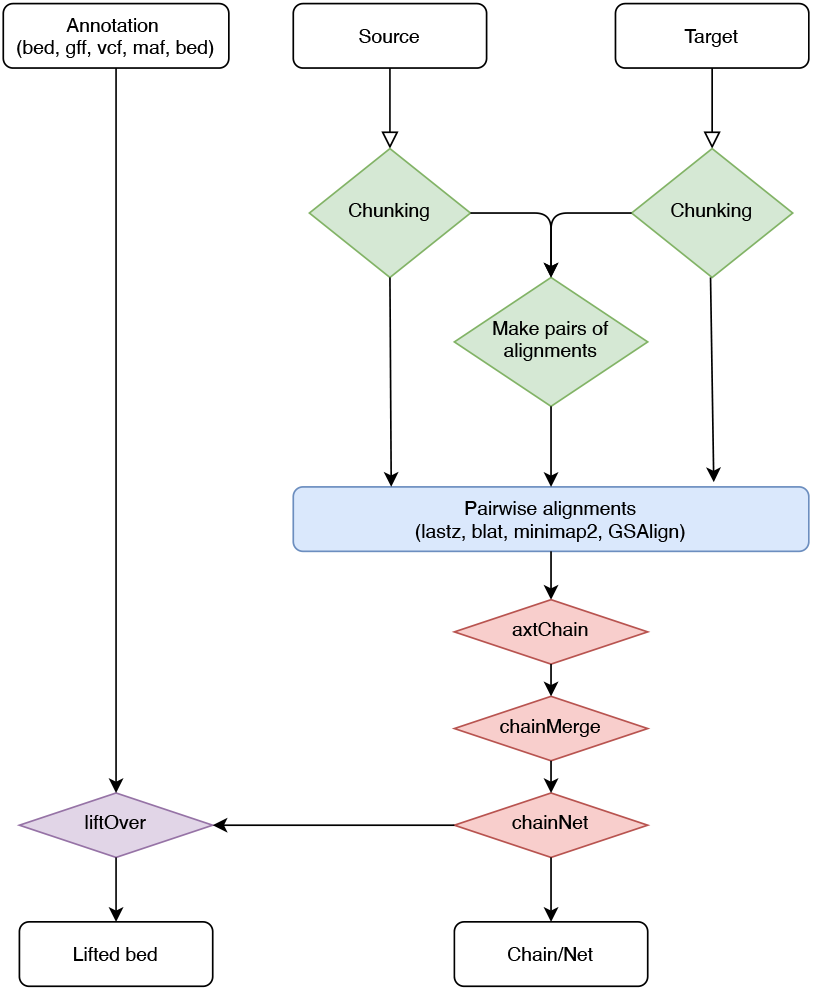
Scheme of the workflow of nf-LO with the chunking (step 1, in green), alignment (step 2, in blue), generation of the liftover files (step 3, in red) and optionally lifting of the variants to the target genome (step 4, in purple).

The alignment phase can be performed in different ways, depending on the type and sensitivity required by the user. For same-species alignments, we provide native support for both blat (Kent 2002), the aligner of choice for same species liftover files from the UCSC genome browser, and GSAlign (Lin and Hsu 2020), a new, high speed same-species alignment software. For performing different-species liftovers, nf-LO also incorporates lastz (Harris 2007), used by the UCSC genome browser to generate between species liftover files, and minimap2 (Li 2018), one of the fastest genome-to-genome aligners. All these aligners are integrated within the workflow, keeping unchanged the UCSC backbone for downstream stages (UCSC 2018). We provide canned configurations for each aligner based on how distant the two genomes are (e.g. near or far), with the possibility to provide sets of custom parameters to achieve the desired balance between speed and sensitivity (Supplementary table 1). nf-LO achieves similar liftover coverage as liftover files from UCSC with appropriate tuning of the parameters (Supplementary table 2).

The third stage processes the alignments analogously to the UCSC processing pipeline, obtaining the chain-net files to perform the actual liftover. Finally, the fourth step supports both the standard bed format with the LiftOver software, or several additional formats using CrossMap (Zhao et al. 2014), including popular formats such as VCF, BAM and GFF.

In conclusion, we provide a transposition of the UCSC liftover pipeline within the Nextflow language, together with the necessary containers to run the analyses, allowing an easy, streamlined implementation in any Unix-based system. We believe that this workflow will be of use across genomics studies, facilitating research work and enabling data interpretation.

## Code availability

The code described in the paper is publicly available on GitHub at the repository https://github.com/evotools/nf-LO. The documentation for the software can be accessed in the wiki page of the website (https://github.com/evotools/nf-LO/wiki).

## Authors’ contributions

AT and JP conceived the study. AT developed the software. AT and JP tested the code. AT and JP contributed to data interpretation and drafted the manuscript. All authors reviewed and approved the final manuscript.

## Acknowledgements

This work was supported by BBSRC grants BB/T019468/1 and BBS/E/D/10002070.

## Captions

Supplementary Table 1 – Comparison of the run times of different aligners and configurations using the human genome GRCh38 as the source and four other large genomes (>1Gbp) as targets on a Scientific Linux 6.9 system with AMD Opteron 6376 2.3GHz 64-cores and 500 GB of RAM.

Supplementary Table 2 – Coverage for the liftover chain files both generated by us and those available from the UCSC genome database, calculated by converting the chain files to maf (chainToAxt > axtToMaf) and then using mafCoverage (Earl et al. 2014).

## Notes

https://github.com/evotools/nf-LO

### Competing Interest Statement

The authors have declared no competing interest.

